# The seasonal dynamics of bud dormancy in grapevine suggest a regulated checkpoint prior to acclimation

**DOI:** 10.1101/2021.02.09.430520

**Authors:** Yazhini Velappan, Tinashe G Chabikwa, John A Considine, Patricia Agudelo-Romero, Christine H Foyer, Santiago Signorelli, Michael J Considine

## Abstract

Grapevine (*Vitis vinifera* L.) displays wide plasticity to climate and seasonality, ranging from strongly deciduous to evergreen. Understanding the physiology of decisions to grow or quiesce is critical for improved crop management, prediction, and the adaptability of production to alternative climate scenarios. The perenniating bud (N+2) is a major economic unit and focus of study. Here we investigated the physiology and transcriptome of cv. Merlot buds grown in a temperate maritime climate from summer to spring in two consecutive years. The changes in bud respiration, hydration and internal tissue oxygen data were consistent with the transcriptome data. ABA-responsive gene processes prevailed upon the transition to a deep metabolic and cellular quiescence in the bud during autumn. Light, together with hypoxia and redox signalling presided over the resumption of nuclear and cellular growth in the transition to spring. Comparisons with transcriptome data from bud burst studies revealed a number of regulatory candidates for the orderly resumption of growth in spring, including components that may integrate light and temperature signalling. Importantly however, the bud burst forcing data, which is widely used as a measure of bud dormancy, were not consistent with the physiological and transcription data. We hypothesise the existence of a physiological checkpoint following bud set in summer, which if not met results in extreme quiescence. Collectively this is the most integrated developmental dataset of the latent bud of cultivated grapevine, and establishes a platform for systems approaches to study seasonal plasticity.

**One sentence summary:** Physiology and transcriptome data provide strong evidence of a regulatory checkpoint prior to acclimation and dormancy in latent grapevine buds.

## INTRODUCTION

The dormancy of the latent or perenniating bud is a seasonally entrained condition that is expressed by many perennial plant species. Improved knowledge of how seasonal cues entrain bud development and dormancy is important in order to manage and mitigate the effects of regional and global climate change in perennial forest and crop systems. Despite recent advances in understanding the seasonality of bud dormancy in some temperate and boreal species (reviewed by Rohde and Bhalerao, 2007; Tanino et al., 2010; Cooke et al., 2012; van der Schoot et al., 2013; Singh et al., 2017), our knowledge of the regulation of bud dormancy in cultivated grapevine (*Vitis vinifera* L.) remains poor. There is a considerable gap in our understanding of the onset and depth of bud dormancy in grapevine.

Dormancy is defined as the failure of a quiescent but viable and intact meristem to resume growth in a permissive environment (Rohde and Bhalerao, 2007). Photoperiod and temperature are the primary ecological cues driving the onset of, and release from dormancy. However, dormancy is considered to be a quantitative condition; ecological cues and developmental state influence the timing and depth of dormancy (Cooke et al., 2012). Genetic analyses in some woody perennial species have dissected the regulation of correlated developmental processes, such as the cessation of growth, bud set, and the onset of dormancy. For example, ethylene insensitive birch (*Betula pendula*), expressing a dominant negative ETR1 ethylene receptor (*etr1-1*), did cease shoot growth but failed to set terminal buds during the transition to short days (Ruonala et al., 2006). Although delayed, the axillary buds did enter dormancy. In wild type *Vitis* spp., the transition to short days accelerated the cessation of shoot growth, bud set, periderm formation and the onset of dormancy (Fennell and Hoover, 1991; Wake and Fennell, 2000; Grant et al., 2013). Short day treatment also enhanced the depth of dormancy (Fennell and Hoover, 1991).

Data on the depth of dormancy in *Vitis* spp. are considerably diverse. The depth of bud dormancy is typically measured in a bioassay of single node explants, grown in forcing conditions (Lavee and May, 1997; Camargo Alvarez et al., 2018). However, dormancy may be calculated as a percentage of buds burst within a given period (e.g. 28 d), or as the time required to reach 50 % bud burst (BB_50_). The two metrics may give quite different seasonal profiles in the depth of dormancy. Measurements of several grapevine varieties from 26° to 34° latitude suggest that the depth of dormancy increases to a maximum prior to, or during early winter (Lavee and May, 1997; Parada et al., 2016; Rubio et al., 2016; Zheng et al., 2018a). A selection of studies illustrate a distinct behaviour, where the depth of dormancy shows a pronounced peak in late summer before declining prior to winter (Pouget, 1963; Nigond, 1967; Cragin, 2015). Additionally, the depth of dormancy varies widely. Measured as BB_50_, Rubio et al. (2019 and references therein) showed a range in peak dormancy for cv. Thompson Seedless of *ca*. 30 – 45 d in two climate regions (33°34’S latitude *cf*. 30°02’S), while earlier studies of cv. Merlot (44°50’N; Pouget, 1963a) and cv. Carignan (43°36’N; Nigond, 1967) showed BB_50_ of over 200 d. Considerable intra-annual and inter-climate plasticity has been shown in other perennial fruit trees (El Yaacoubi et al., 2016).

Against a background of a growing number of extensive molecular and biochemical studies of the later stages of bud development in grapevine (Halaly et al., 2008; Ophir et al., 2009; Meitha et al., 2015; Zheng et al., 2015; Sudawan et al., 2016; Khalil-Ur-Rehman et al., 2017; Meitha et al., 2017; Signorelli et al., 2018; Zheng et al., 2018a), few have followed development for an extended period of time, from the onset of dormancy. The most extensive catalogue to date spanned the transcriptional state of the bud from shoot development, prior to the summer solstice and periderm formation, through to bud burst following winter (cv. Tempranillo, Madrid, Spain, 40°28’N; Diaz-Riquelme et al., 2012). Pronounced circannual rhythms were seen in homologues of flowering pathway transcriptional regulators as well as major functional categories of genes, e.g. photosynthesis and regulation of the cell cycle (Diaz-Riquelme et al., 2012). No physiological data or experimental manipulation were presented, however. Other more acute temporal studies have demonstrated the role of photoperiod in floral initiation (Sreekantan et al., 2010), cell wall thickness (Rubio et al., 2016) and dormancy onset (Fennell et al., 2015).

Here, we have established for the first time an integrated platform of physiological and transcriptome data to investigate the seasonal signalling relationships that govern the quiescence states of the latent bud in cultivated grapevine. The physiological data were consistent with the dynamic observations of BB_50_ by Pouget (1963) in cv. Merlot in Bordeaux, France, revealing an extreme resistance of explants to resume growth when sampled in late summer. This resistance however, was not consistent with the respiration, hydration or internal tissue oxygen data of the bud, nor the prevailing influences at the level of the transcriptome. These indicated that metabolic and gene regulatory activity was highly primed during summer, followed by an increasingly quiescent state during autumn and the onset of winter. We propose the existence of an acclimation checkpoint in late summer, which if not met will trigger a stress-induced dormancy akin to secondary dormancy of seeds.

## RESULTS

### The within year dynamics of dormancy and physiology of grapevine buds

We evaluated the competence of bud growth of cv. Merlot explants sampled from mid-summer through to early spring over two consecutive years (Margaret River, Australia; 33°47’S, 115°02’E). Single node cuttings from a commercial vineyard were transferred to a controlled temperature growth room for up to 350 d (forcing conditions). The first sample point was in late January (*ca*. 4 weeks after the summer solstice at 22^nd^ December). The data revealed a pronounced but transient peak in bud dormancy (BB_50_), during the period from late January to April, reaching a climax during February/March at BB_50_ *ca*. 250 d (Figure 1b). The BB_50_ declined at a similar rate as it developed, from late March to early April, from BB_50_ *ca*. 240 to 70 d. Thereafter the BB_50_ continued to decline at a more linear but steady rate towards August/September, at the beginning of spring (Figure 1b). Between 70-80 % buds sampled during the summer/autumn climax period burst within 350 d and the remaining were necrotic. Positive control buds treated with hydrogen cyanamide (H_2_CN_2_) reached >95 % bud burst (Figure 1b, refer below), indicating that the necrotic buds were viable at the time of sampling. Notwithstanding, the buds sampled during the February/March period were remarkably resilient. These trends were consistent in both 2015 and 2016 (Supplemental Table S1), and consistent with the patterns observed in cv. Merlot and cv. Carignan in France (Pouget, 1963; Nigond, 1967).

**Figure 1.**
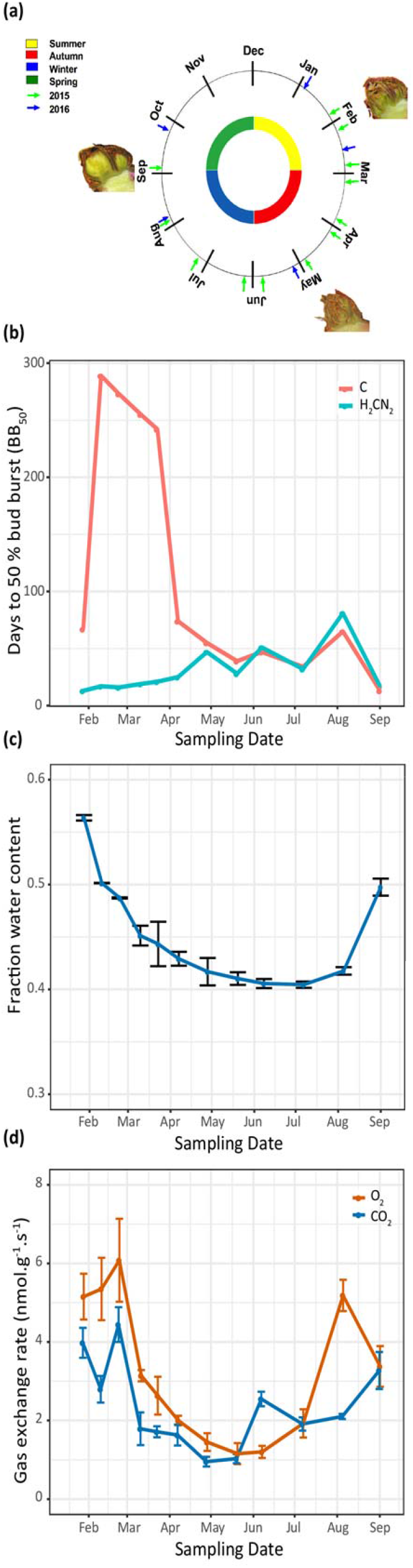
Dormancy and physiology of grapevine buds sampled throughout the season in the Margaret River region of Western Australia (2015, 33°S latitude). **(a)** Phenological calendar showing the sampling dates for 2015 and 2016 in this study and corresponding seasons. The calendar position of the bud images indicates the sample times used for RNA sequencing. **(b)** The depth of dormancy of single-node explants collected in 2015 and grown forcing conditions, expressed as the time to 50 % bud burst (BB_50_; n=50). Control (C) and buds treated with 0.31 M hydrogen cyanamide (H_2_CN_2_). **(c)** Water content (n=3) and **(d)** respiration (n=4) of samples collected at the corresponding dates. Vertical bars on **(c)** and **(d)** represent *s*_*x*_. Data for 2016 provided in Supplemental Table S1.

The application of H_2_CN_2_ dramatically accelerated the rate of bud burst (BB_50_) when applied to explants sampled during the period from February through to April (Figure 1b; Supplemental Table S1). Thereafter, H_2_CN_2_ had very little effect on the rate of bud burst. The H_2_CN_2_-treated buds did however show an interesting developmental trend; a moderate but progressive increase in the BB_50_ from January (mid-summer) through to August (end winter), from BB_50_ *ca*. 30 to 70 d, before an abrupt decline *ca*. 20 d in September, which was shortly before the time of natural bud burst in the field (Figure 1b). The trend was observed in both the 2015 and 2016 seasons (Figure 1b; Supplemental Table S1).

Matching bud material was used to determine hydration levels, respiration rates and internal tissue oxygen status. The hydration levels of buds sampled in late January (mid-summer) was *ca*. 55 gH_2_O.100g FW^-1^ (Figure 1c, Supplemental Table S1). Hydration levels initially declined quite rapidly until April, then more gradually towards a point of inflection at 40 gH_2_O.100g FW^-1^ by July, and increased thereafter. The O_2_ consumption and CO_2_ production rates remained high until the end of February, before declining rapidly and in parallel by *ca*. 2-3-fold by April. The gas exchange rates then declined more gradually to reach a minimum during May/June, and increased thereafter (Figure 1d, Supplemental Table S1). We expected that changes in respiration and hydration may also influence the pO_2_ within buds. Figure 2 shows a moderate but consistent decline in the pO_2_ at the meristematic region of the bud (*ca*. 1500-2000 μm) from February through to August, before returning to a more normoxic state by September.

**Figure 2.**
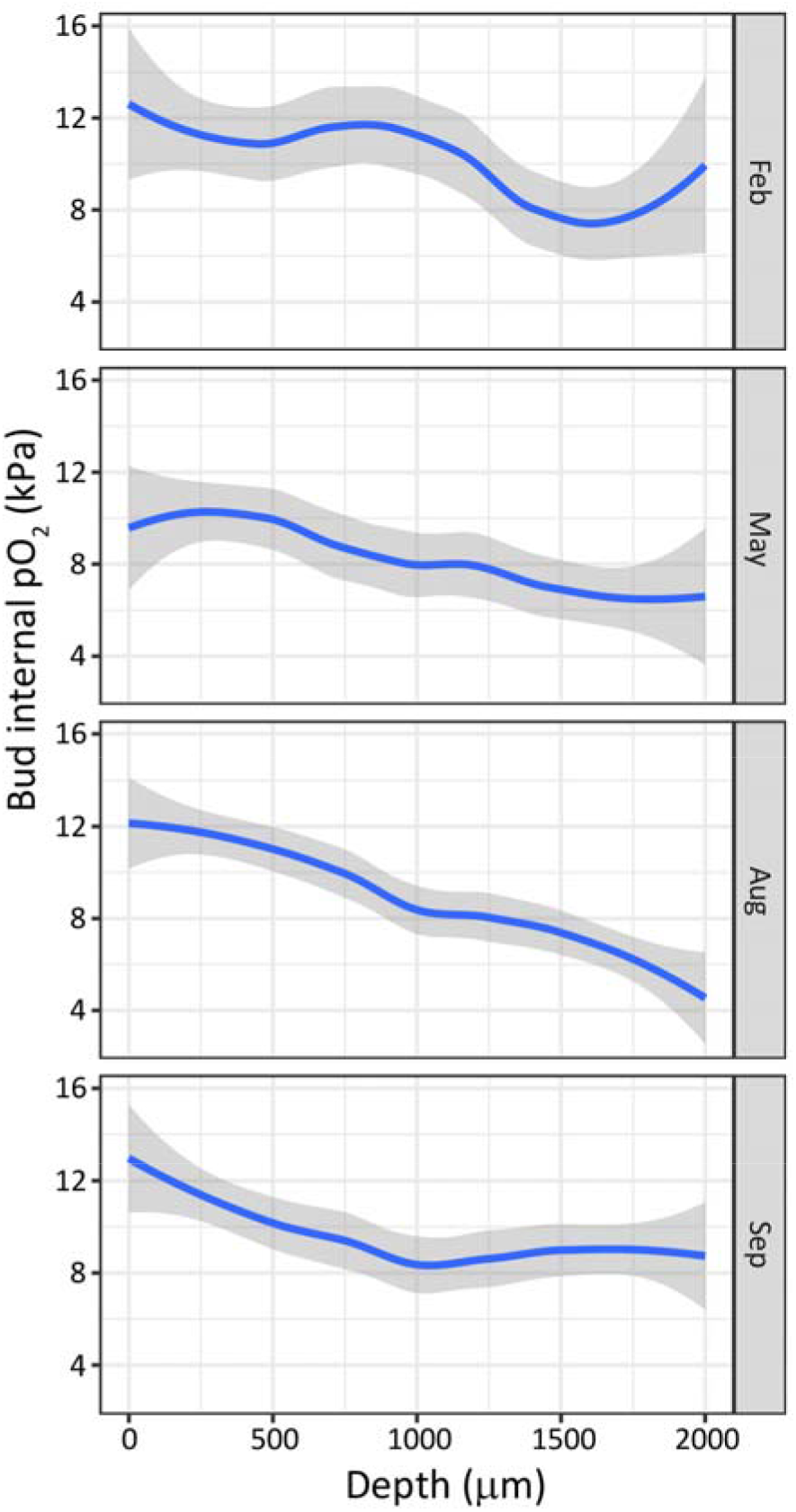
Tissue oxygen partial pressure (pO_2_) profiles in grapevine buds sampled from summer to spring in the Margaret River region of Western Australia in 2015 and 2016. Buds were collected over two consecutive seasons (2015, 2016), n=6 individual buds, with the exception of February (2016 only, n=3). Sample dates (Feb, May, Aug and Sep) are shown in Figure 1a. Data show the pO2 from immediately within the outer scales (0 μm depth) to the region of the meristematic core of the primary bud (2000 μm). Atmospheric oxygen is *ca*. 21 kPa pO_2_. The plot represents a regression curve with 95 % confidence intervals.

Taken together, the data show a remarkable seasonal dynamic of BB_50_, with a pronounced but transient peak during late summer/early autumn. This behaviour was not reflected by changes in respiration or hydration levels of the buds. Hydration levels began to decline before changes in BB_50_ and respiration. The decline in respiration also appeared to precede the decline in BB_50_ in autumn, however the rate at which BB_50_ declined was more rapid. The respiration rate then began to increase in early winter, prior to the increase in hydration levels. Natural bud burst occurs in early to mid-spring, and there was an abrupt increase in hydration at this point. Internal tissue oxygen status was less informative, although there appeared to be a major transition in August, with oxygen levels at the core of the bud becoming more hypoxic, at the time when respiratory oxygen consumption reached a peak prior to natural bud burst in September (Figure 2).

### Relationship of dormancy and physiology to climate indices

Site weather data were obtained for the two years of the study (Supplemental Figure S1, Supplemental Table S1). The maximum daylength was *ca*. 14 hrs at the summer solstice, which had declined by *ca*. 1 hr by the earliest sample point 4 weeks later in late January. The minimum daylength was *ca*. 10 hrs at the winter solstice. Temperature maxima/minima ranged from *ca*. 28/12 °C in January to *ca*. 16/5 °C in July. Rainfall data were also collected, showing a temperate pattern of rain falling predominantly during the winter months (Supplemental Table S1). While the study vineyard site was not irrigated, there was 78 mm and 170 mm rain in the January to March period of 2015 and 2016, indicating adequate water availability during the drier summer months.

The accumulated chilling in the vineyard was calculated according to three commonly used models (Supplemental Figure S1). Chilling began to accumulate from April onwards (*ca*. 100 d post-solstice), at a near-linear rate towards and beyond the time of natural bud burst in the field in September (*ca*. 270 d post-solstice). The chill summation was similar for both Utah models (> 1000 units) and *ca*. 260 units according to the base 7.2 °C calculation (Supplemental Figure S1c).

### Differential transcriptome analysis

RNA extracted from buds collected at six time points in 2015 were sequenced. After pre-processing, normalized log2 counts per million (CPM) data were used to generate a Principal Component Analysis (PCA) plot, showing that replicates of each time point largely clustered together, and that the September (S9) samples formed the most distinct cluster (Figure 3a). The first component (32.36%) seemed to be largely affected by developmental age, while the second component (26.81 %) appeared to be linked to metabolic activity and degree of hydration (Figure 1,Figure 3a). A similar discrimination along developmental trends was reported by Diaz-Riquelme et al. (2012). Differential expression analysis showed very few differentially expressed genes (DEGs) between January-February and February-March time-points while PCA showed February and March clustered together (Figure 3a). Considering this, and guided by the BB_50_, respiration, hydration and chilling data (Figure 1, Supplemental Figure S1), we chose to refine the comparisons to three:

**Figure 3.**
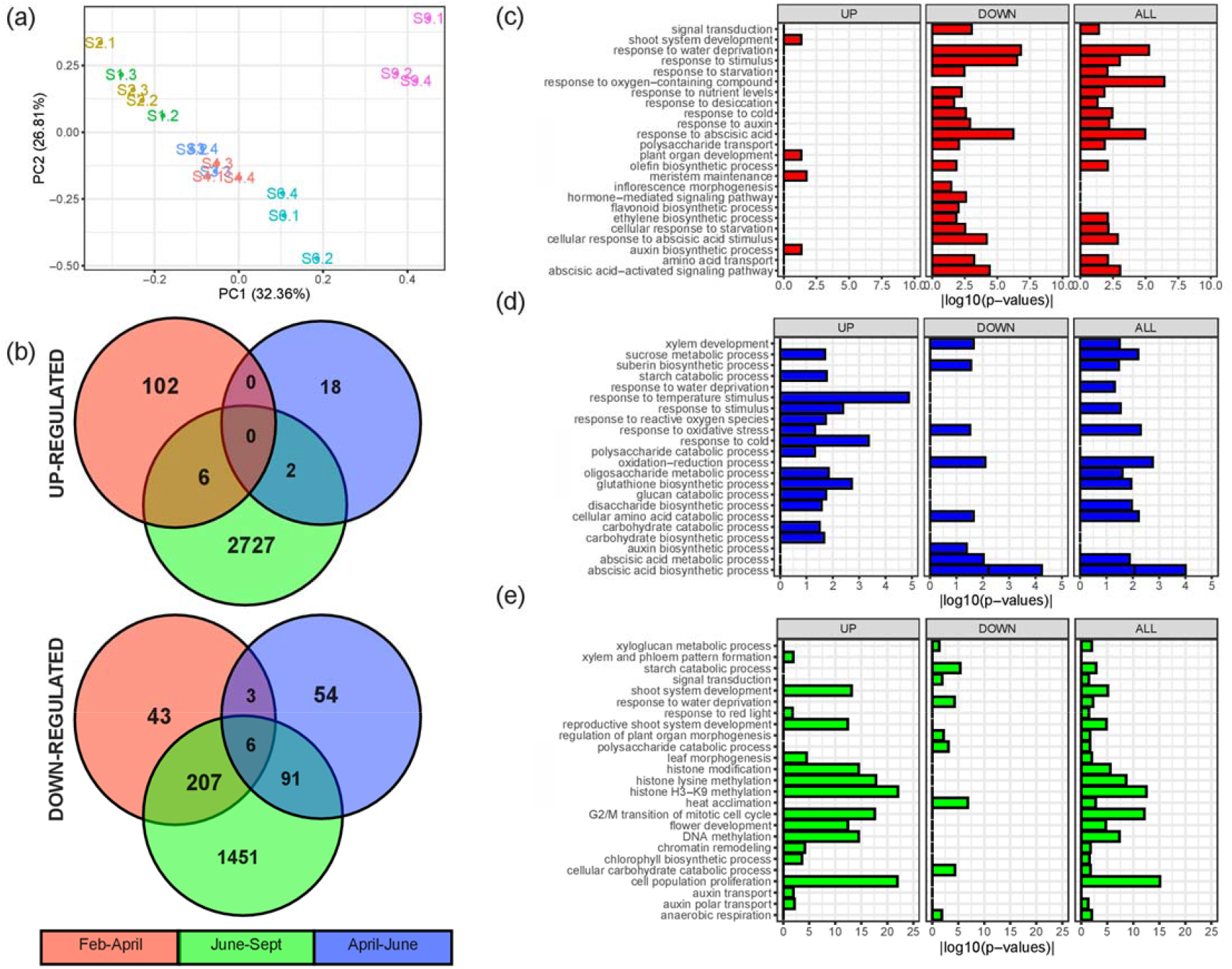
PCA plot of normalized count data, and Venn diagrams and functional enrichment analysis of differentially expressed genes (DEG) from grapevine buds collected at different stages of dormancy transition February to April, April to June and June to September. **(a)** Principal components analysis (PCA) of normalized read counts (S1-January; S2-February; S3-March; S4-April; S6-June; S9-September). **(b)** Venn diagrams indicating the number of significant (FDR ≤ 0.01, Log_2_FC |1|) DEGs across three comparisons and the overlap between each set of genes separated into up- and down-regulated genes. Horizontal bar plots of selected functional categories significantly enriched (*P* ≤ 0.05 using the hypergeometric test) in the developmental comparisons February-April **(c)**, April-June **(d)** and June to September **(e)**. Three different functional enrichment analyses were performed for each comparison, where “ALL” (up-plus down-regulated genes), “UP” (up-regulated genes) and “DOWN” (down-regulated genes) make reference to the list of genes assessed.

i. April/February (*ca*. 95/40 d post-solstice), representing rapid decline from the dormancy peak and a parallel decline in respiration and hydration.
ii. June/April (*ca*. 175/95 d post-solstice), representing the period prior to winter, accompanied by the minimum hydration and respiration rates, and the onset of chilling accumulation.
iii. September/June (*ca*. 240/175 d post-solstice), representing the period during winter, accompanied by the resumption of hydration and respiration and the majority of chilling accumulation.

Following differential expression analysis, annotations were assigned to the core set of DEGs against the *V. vinifera* 12X v.2.1 annotation file, and functional enrichment was performed based on gene ontology (GO) analysis.

A total of 4490 DEG were identified across the three stages of dormancy transition (Supplemental Table S3), which represents *ca*. 10 % of the predicted genes in the *V. vinifera* genome (Canaguier et al., 2017). A Venn diagram of DEGs revealed that the June-September transition was the most discriminating, as over half of all DEGs (2727) were uniquely up-regulated and one third (1451) uniquely down-regulated during this transition (Figure 3b). Few genes were consistently up-regulated between successive transitions; no genes were consistently up-regulated throughout the experiment and only 6 genes consistently down-regulated (Figure 3b). These data suggest stage-specific gene expression profiles during the subsequent developmental transitions. Functional enrichment analysis showed that genes involved in hormone signalling, nutrition, cell proliferation, morphological development, epigenetic regulation were highly regulated during the time course (Figure 3c-e). Comprehensive gene ontology (GO) enrichment results are presented in Supplemental Table S4.

### The transition from summer to autumn

The April/February transition represented a 4-fold decline in BB_50_ and parallel declines in hydration and cellular respiration (Figure 1b-d). The GO enrichment, together with the DEG data revealed a pronounced decline in the expression of response to stimuli functions, including abiotic, hormone and nutrient stimuli (Figure 3c; Supplemental Table S3, S4). Expression profiles and unique identifiers of all genes subsequently described are shown in Supplemental Table S5.

Down-regulated abiotic response genes included four homologues of *GALACTINOL SYNTHASE* as well as a *RAFFINOSE SYNTHASE* and *STACHYOSE SYNTHASE*, which collectively encode functions to synthesise raffinose family oligosaccharides (RFO; Supplemental Table S5). The *VACUOLAR INVERTASE 2* (*VI2*), involved in sucrose catabolism, a sugar transporter *SWEET17* (previously known as *NODULIN MtN3*), and *TREHALOSE-6-PHOSPHATE PHOSPHATASE A* (*TPPA*) were down-regulated, as were an *AMMONIUM TRANSPORTER 2* and an *AMINO ACID PERMEASE*, indicative of a transcriptional down-regulation of nutrient transport functions from February to April. Genes involved in abscisic acid (ABA) biosynthesis and signalling were down-regulated; two *9-CIS-EPOXYCAROTENOID DIOXYGENASES*, and *MOTHER OF FT AND TFL1* (*MFT*), together with four *LATE EMBRYOGENESIS ABUNDANT* and two homologues of *SENESCENCE-ASSOCIATED GENE 101*. Genes encoding functions that generate or process reactive oxygen species (ROS) were also down-regulated during this transition, including a *1-CYSTEINE PEROXIREDOXIN*, two *PEROXIDASES* and a *RESPIRATORY BURST OXIDASE HOMOLOGUE (RBOH)*.

Against this, a small number of functions assigned to phyllome development were significantly enriched in the GO data, and up-regulated during this transition (Supplemental Table S4). For example, a *DORNROSCHEN-like* (*DRN-LIKE*/*ESR2*) ethylene response factor (ERF) transcription factor, two *GROWTH-REGULATING FACTOR* genes, and two gibberellic acid (GA)-responsive genes were up-regulated (Supplemental Table S5). Interestingly *DRN-LIKE* is the most homologous grapevine gene to the poplar *EARLY BUD-BREAK 1* (*EBB1*), which is a positive regulator of bud burst (Yordanov et al., 2014; Busov et al., 2016). Taken together, these data indicate that the February condition, where the time to BB_50_ was maximal, was highly regulated, with relatively high expression of genes functioning in responses to abiotic stimuli, ABA and ROS. The trend to a decline in ABA synthesis and responses may however indicate that ABA levels remained high and were in the process of down-regulation by negative feedback.

### The transition from autumn to winter

By comparison with the previous transition, April to June represented quite modest declines in the BB_50_, hydration and the rate of respiration (Figure 1). The number of genes differentially regulated was also the least of each of the transitions, and the majority were down-regulated (Figure 3b, Supplemental Table S3). Nevertheless, GO analysis revealed a prominent enrichment of carbohydrate metabolism, namely sucrose metabolism, starch catabolism, polysaccharide catabolism, oligosaccharide metabolism, glucan catabolism, disaccharide biosynthesis, carbohydrate catabolism and biosynthesis. None of these GO were enriched in the other two transitions (Figure 3), suggesting a specific regulation of carbohydrate metabolism during the autumn to winter transition. GO analysis also revealed that response to temperature, cold and oxidative stress/ROS were more enriched in June, relative to April (Figure 3d; Supplemental Table S4).

Transcripts coding for ABA synthetic enzymes; *NCED6* and *ABA DEFICIENT 2* (*ABA2*) were down-regulated, as well as genes coding for the synthesis of RFOs, which were down-regulated in the previous transition (Supplemental Table S3, S5). A *PEROXIDASE 1* and *GLUTATHIONE PEROXIDASE 8* were representative of declines in ROS processing functions, although a *ROXY1* thioredoxin superfamily protein was up-regulated. The *γ–GLUTAMYLCYSTEINE SYNTHETASE* (*ECS1*), which codes for the first committed step of glutathione synthesis was also up-regulated. Up-regulation of temperature-regulated genes were represented by two homologues of the *COR27* cold regulated gene. Together, these are consistent with acclimation to abiotic stress and desiccation, and a more metabolically quiescent state than the preceding phase.

### The transition from winter to spring

The final developmental comparison of September/June was accompanied by a relatively modest decline in the BB_50_, however hydration, the rate of respiration and tissue oxygen status increased markedly, as did photoperiod and the cumulative exposure to chilling (Figures 1-2, Supplemental Figure S1). By this stage buds had not burst naturally in the field, although bud burst was imminent. The number of uniquely up-or down-regulated genes was the greatest of any transition (Figure 3b). The GO analysis showed a strong enrichment of functions assigned to DNA replication and epigenetic modification in this transition, such as histone modification, histone lysine methylation, histone H3-K9 methylation, G2/M transition of mitotic cells, DNA methylation and chromatin remodelling (Figure 3e; Supplemental Table S4). In addition, GO analysis revealed a significant enrichment of transcript involved in shoot and vascular system development. Numerous functions were unique or regulated in an opposite direction to changes of previous transitions.

Four homologues of *CHROMOMETHYLASE 2* and *3*, a *METHYLTRANSFERASE 1* (*MET1*) and a chromatin remodelling *DECREASE DNA METHYLATION 1* (*DDM1*) were up-regulated in September (Supplemental Table S5). In addition, *ARABIDOPSIS TRITHORAX-RELATED PROTEIN* (*ATXR*) *5* and *ATXR6*, two genes encoding a nucleolar histone methyltransferase-related protein, and a histone deacetylase were up-regulated, as were numerous genes coding for histone subunits. *VERNALIZATION 1* (*VRN1*), *VERNALIZATION 3-LIKE* (*VEL1*) and other related epigenetic factors are also outlined below, while cell cycle genes are outlined in a section below.

ABA- and stress-responsive genes were widely down-regulated. These included homologues of *NCED4, LEA4-5*, dehydrin *XERO1*, three cold regulated *COR27*, and four PP2C genes; *ABA INSENSITIVE 1* (*ABI1*), *ABA-HYPERSENSITIVE GERMINATION 3* (*AHG3*) and two *HIGHLY ABA-INDUCED* genes (*HAI2* and *HAI3*; Supplemental Table S5). Two of the *COR27* were previously up-regulated from April to June. Homologues of SNF1-related protein kinase genes were down-regulated (*SNRK2*.*6, SNRK3*.*14, SNRK3*.*8, AKINBETA 1, KING1*), as were genes coding for dormancy associated proteins DELAY OF GERMINATION 1 (DOG1) and DORMANCY ASSOCIATED PROTEIN 1 (DRM1). Against this, genes encoding auxin transport and signalling functions were predominantly up-regulated. These included two homologues encoding the auxin efflux carrier *PIN-FORMED 1* (*PIN1*), plus *PIN2, PIN5* and *PIN6*, together with the auxin receptor *TRANSPORT INHIBITOR RESPONSE 1* (*TIR1*) and number of auxin responsive factor proteins. Cytokinin signalling genes were also up-regulated, including a number of homologues of *ARABIDOPSIS RESPONSE REGULATOR* family genes. Meanwhile GA- and ethylene-related functions were differentially regulated. GA biosynthetic genes were up-regulated, as was a homologue of the DELLA-degrading F-box protein *SLEEPY2*, while a GA receptor (*GID1B*) and two genes of GASA domain proteins were down-regulated. Similarly, the ethylene biosynthetic gene *ACC OXIDASE 1* was up-regulated while *ETR1* and *EIN3* were down-regulated. Together these indicate a finely controlled transition of hormone synthesis and processing during bud development between June and September, with a considerable decline in the influence of ABA against an increase in that of auxin and cytokinin-dependent functions.

Together with the down-regulation of cold-regulated *COR27*, the up-regulation of *VRN1, VEL1* and *DROUGHT SENSITIVE 1* and changes in metabolic functions were consistent with acclimation following stress (Supplemental Table S5). A number of *AMYLASE* genes were down-regulated, while three homologues of *PHOSPHOENOLPYRUVATE CARBOXYKINASE 1*, three *TREHALOSE-6-PHOSPHATE SYNTHASEs* (*TPS*), a *TREHALOSE-6-PHOSPHATE PHOSPHATASE*, three *SUCROSE SYNTHASE* (*SUSY*) and *SEED IMBIBITION 1* and *2*, which encode raffinose synthases were up-regulated. Important genes involved in facilitating transmembrane sugar and nitrogen transport were up-regulated. These included *CELL WALL INVERTASE 1, SWEET 17* and *ERD SIX-LIKE 1*, a homologue of a stress-inducible monosaccharide transporter. Two homologues of *AMINO ACID PERMEASE 2* and a nitrate transporter *NRT1:2* were up-regulated. These reflect a transcriptional up-regulation of gluconeogenesis, sucrose, trehalose and raffinose metabolism and transport functions during this transition.

Cell wall, pectin and cellulose metabolism was widely regulated during this transition, through up-regulation of a number of genes encoding cellulose and glucan synthases pectin methylesterase and invertase/pectin methylesterase inhibitors (Supplemental Table S5).

Genes encoding redox-related functions were largely up-regulated, including homologues of *RBOHD, ROXY1* and *ROXY2* (Supplemental Table S5). Ascorbate synthesis was positively regulated through *L-GALACTONO-1,4-LACTONE DEHYDROGENASE* (*GLDH*), as was cysteine through *Ο-ACETYLSERINE (THIOL) LYASE* (*OAS-TL*), while glutathione synthesis was mildly down-regulated (*ECS1*). Response to hypoxia was also evident; homologues of 10/49 conserved hypoxia response genes were regulated only at this final transition (Mustroph et al., 2009; Supplemental Table S5). These included *ACC OXIDASE 1, LOB DOMAIN-CONTAINING PROTEIN 41* (*LBD41*), *SUSY4, PLANT CYSTEINE OXIDASE 1* (*PCO1*), and a homologue of *RBOHD* (mentioned above), which were strongly up-regulated, consistent with the function in hypoxic response. Also up-regulated were *RELATED TO AP 2*.*3* (*RAP2*.*3*) and *LITTLE ZIPPER 2* (*ZPR2*), which have demonstrated functions in hypoxia (Weits et al., 2019).

Numerous genes encoding cell identity, meristem and flowering functions were up-regulated in addition to those mentioned above (*VRN1, VEL3, ZPR2, LBD41;* Supplemental Table S5). These include multiple homologues of *WUSCHEL-RELATED HOMEOBOX* proteins (*WOX1, 3, 4, 9*), *CLAVATA 1* (*CLV1*), *CLAVATA3/ESR-RELATED 44* (*CLE44*) *and SHOOT MERISTEMLESS* (*STM*).

The broad functions represented in the transcriptional changes from June to September are consistent with a post-acclimation reorganisation of nuclear, metabolic and cellular structure and function. The prominence of redox- and hypoxia-responsive genes may reflect a relationship to chromatin regulation and differentiation.

### Comparison of quiescence and bud burst data

In earlier studies we have identified a number of developmentally and light-regulated transcripts during the early transition towards bud burst (Meitha et al., 2017; Signorelli et al., 2018). A comparison of those DEG data sets with the present set revealed 80 genes that were differentially regulated in all studies (Supplemental Table S5). Of these only two were not differentially regulated in the transition from winter to spring here. There was considerable agreement between those genes regulated between winter and spring and those light-regulated in the first 144 h of bud burst. This included the down-regulation of *DRM1* and a SNF1-related protein kinase *KING1*, and up-regulation *CRYPTOCHROME 3 (CRY3)*, glutamyl t-RNA reductase *HEMA1, GENOMES UNCOUPLED 4* (*GUN4*) and *PROTOCHLOROPHYLLIDE OXIDOREDUCTASE C* (*PORC*).

### *Cis*-regulatory element enrichment identifies common and unique stress- and developmental gene regulation

To gain further insight to the prevailing transcriptional control during dormancy transitions, we carried out enrichment analysis for known and *de novo cis-*regulatory motifs in the 1.2*-*kb upstream region of the DEGs (Figure 4; Supplemental Table S6). A remarkably large proportion of the total number of enriched motifs were common to all transitions (Figure 4b). Figure 4a shows the most significant (lowest p-value) motifs identified, and Figure 4c shows a selection of the unique motifs identified at each transition. The TEOSINTE BRANCHED1, CYCLOIDEA, PCF (TCP) and ETHYLENE-RESPONSIVE TRANSCRIPTION FACTORS (ERF) binding sites featured prominently throughout (Figure 4a). Two WRKY motifs were specifically represented in the April/February dataset. Meanwhile the MYB3R and SPL motifs predominated among enriched motifs in the September/June dataset, suggesting a prominent role of the MYB3R and SPL family transcription factors during the final transition preceding bud burst.

**Figure 4.**
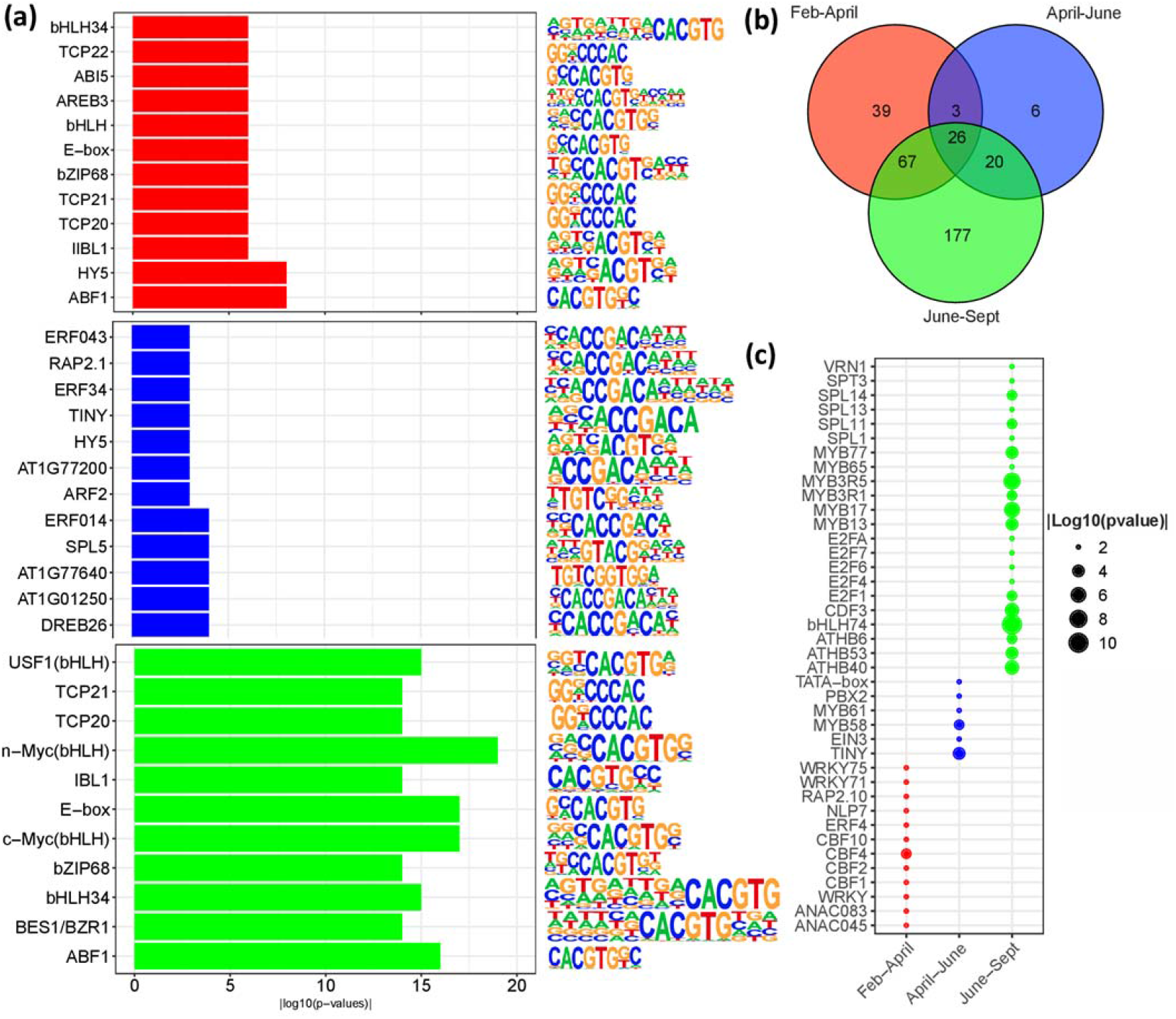
Enrichment analysis of known *cis*-regulatory binding motifs identified in promoter regions DEGs (FDR ≤ 0.01, Log_2_FC |1|) at different stages of bud dormancy transition for the comparisons, February to April, April to June and June to September. **(a)** Bar plots showing the 10 most significantly enriched motifs (lowest *P* value) for the comparisons, February-April (red), April-June (blue) and June to September (green). **(b)** Venn diagram discriminating the common and unique motifs in the three comparisons. **(c)** A scatter plot of selected unique motifs found in each comparison.

### Core cell cycle genes, water-relations and diverse roles of ERF family genes during dormancy transitions

Our earlier studies have indicated oxygen and ethylene signalling (Meitha et al., 2015; Meitha et al., 2017) and cell-cell transport (Signorelli et al., 2020) may hold key regulatory roles in bud development. In addition, we have reviewed the role of cell cycle regulation as a master or slave of dormancy transitions (Velappan et al., 2017). We thus sought to evaluate the transcriptional data here for evidence in support of these functions. Previous studies have identified a set of 61 core cell cycle genes in Arabidopsis (Vandepoele et al., 2002), 138 members of the APETALA2/ethylene-responsive element binding protein (AP2/EREBP) family of plant transcription factors (Riechmann and Meyerowitz, 1998) and 33 aquaporin genes (Ward, 2001) in Arabidopsis. We used *Vitis* 12X v.2.1 annotation file to identify a set of *Vitis* homologues of these genes within our set of DEGs (Supplemental Table S7). We identified 28 cell cycle and 7 aquaporin-related genes (purple and blue panel) which were all up-regulated during the final transition preceding bud burst (June to September; Figure 5, Supplemental Table S7). The dehydration-responsive element binding protein (DREB) subfamily of the AP2/EREBP family were largely down-regulated from June to September (Figure 5). As mentioned above, the grapevine homologue of the poplar *EBB1* (*DRN-LIKE*) was strongly up-regulated in the April/February transition and unchanged thereafter, which is not consistent with the proposed EBB1 function in other species (Yordanov et al., 2014; Busov et al., 2016). Other AP2/EREBP family genes had no definitive expression pattern, which suggests functional diversity amongst this family of transcription factors.

**Figure 5.**
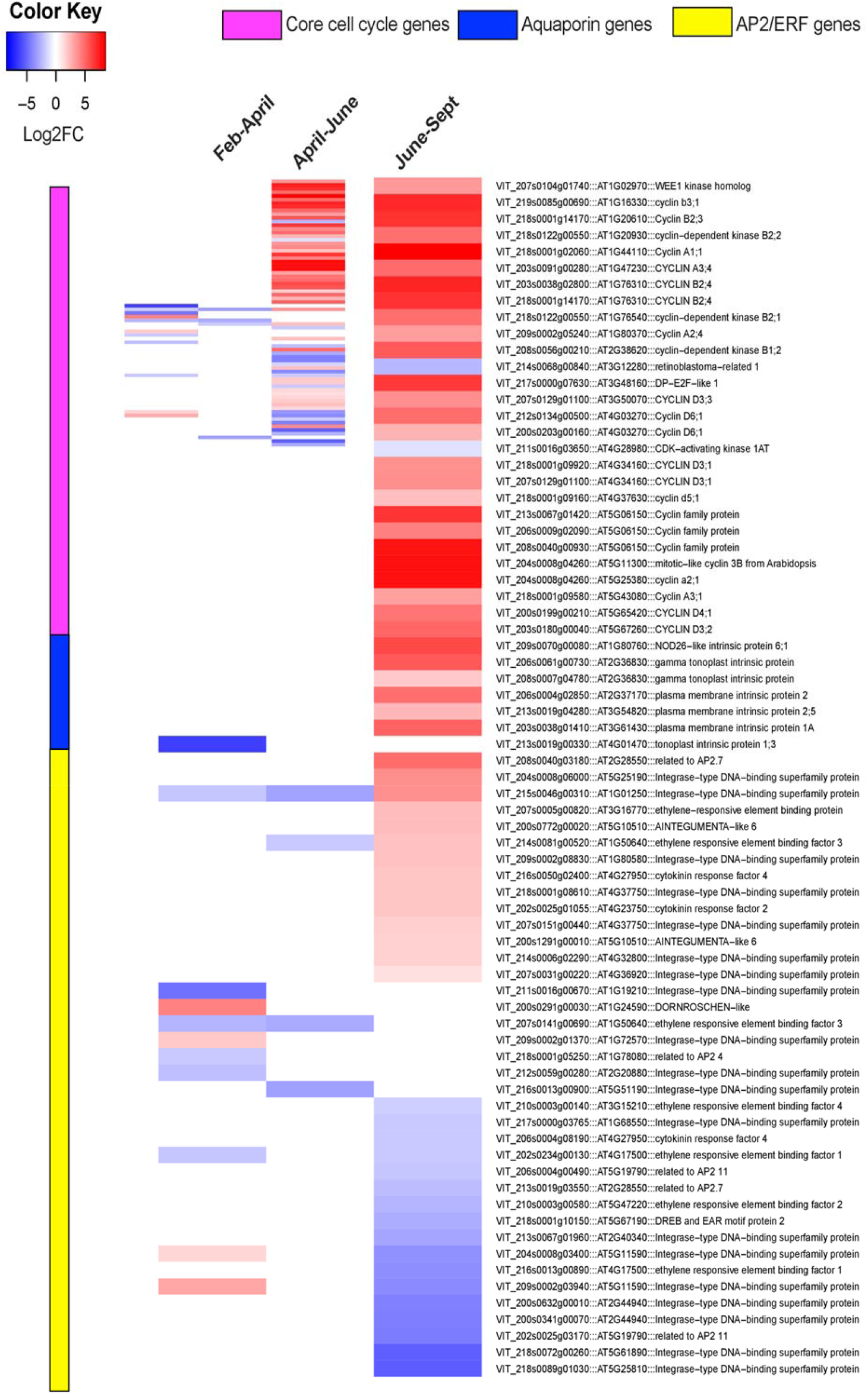
Differential expression of putative cell cycle, aquaporin (water relations) and APETALA2/Ethylene Responsive Factor (AP2/ERF) genes among the DEGs (FDR ≤ 0.01, Log_2_FC |1|) during different stages of bud dormancy transition February to April, April to June and June to September. Homologues of core cell cycle genes of Arabidopsis (Vandepoele *et al*., 2002) are shown in the upper panel of the heatmap (adjacent the purple column). Homologues of The AP2/EREBP family of plant transcription factors of Arabidopsis (Riechmann and Meyerowitz, 1998) are shown in the blue panel of the heatmap. Homologues of aquaporin genes of Arabidopsis (Ward, 2001) are shown in the lower three panels of the heatmap (orange-yellow).

## DISCUSSION

Bud dormancy in woody perennials is understood to be a quantitative condition, however questions remain of the measurable definition of dormancy in this context (Doorenbos, 1953; Samish, 1954; Lang et al., 1987; Rohde and Bhalerao, 2007; Cooke et al., 2012; Considine and Considine, 2016; Camargo Alvarez et al., 2018). For this reason, we predominantly use the terms BB_50_ and quiescence through the following discussion, referring to the phenotype without suggesting the underlying physiological state.

### The extreme quiescence of summer buds is consistent with a checkpoint prior to acclimation

The depth and seasonal dynamics of BB_50_ reported here for cv. Merlot (33°47’S) correlated very well with those reported for cv. Merlot (44°50’N) and cv. Carignan (43°36’N) (Pouget, 1963; Nigond, 1967). There was also reasonable agreement with the seasonal dynamics reported for cv. Thompson Seedless (33°34’S; Parada et al., 2016; Rubio et al., 2016; Rubio et al., 2019), cv. Chardonnay and cv. Cabernet Sauvignon (46°17’N; Camargo Alvarez et al., 2018), although the peak magnitude in days to bud burst reported therein was considerably less. The studies of cv. Thompson Seedless, cv. Chardonnay and cv. Cabernet Sauvignon also reported a more prolonged plateau at the peak of BB_50_ than the studies of cv. Merlot and cv. Carignan (Pouget, 1963; Nigond, 1967). The majority of other bud burst data reported for summer or autumn quiescence of grapevine buds are not comparable to data reported here, for example due to differences in experimental conditions (Or, 2009; Fennell et al., 2015; Pérez and Noriega, 2018).

Trends in respiration data correlated well with that of cv. Merlot (44°50’N; Pouget, 1963) and reasonably well with cv. Thompson Seedless (33°34’S; Parada et al., 2016). Parada et al. (2016) reported an inverse relationship between respiration and BB_50_. Our data show that respiration was not a good predictor of the potential BB_50_, as gas exchange rates of O_2_ and CO_2_ remained high even in buds that developed a BB_50_ >250 d (Figure 1). Water content in our data also failed to explain the dynamics of BB_50_. Water content declined from *ca*. 55 to 45 g H_2_O.100g^-1^ fresh weight from February to March, while respiration and BB_50_ remained high.

The first time point in this study was within 4 weeks of the summer solstice and the first point used for differential gene expression analysis was at *ca*. 7 weeks. We found no differentially expressed genes between these time points. Considering the definition of Rohde and Bhalerao (2007), which we have adopted, the buds developed a highly dormant state over this three week period, despite little change in day length, respiration rate, desiccation, or gene expression, and prior to any exposure to chilling. The physiological and transcriptional changes at subsequent time points were consistent with acclimation to cold and desiccation, and the development of a highly quiescent state of metabolism and genetic control. As a result, we question whether the state of quiescence measured during the transition from late summer to autumn truly reflects bud dormancy *sensu stricto*. We hypothesise that the extreme resistance to resume growth observed in buds sampled in late summer and early autumn was a stress response, akin to secondary dormancy of seeds.

Secondary dormancy is a state induced in non-dormant seeds when exposed to unfavourable conditions (Bewley, 1997; Finch-Savage and Footitt, 2017, and references therein). Seed banks cycle through states of primary and secondary dormancy in nature, as an adaptive response to changes in soil moisture and temperature and light availability. The entry into secondary dormancy of seed appears to be regulated by the balance of ABA and GA signalling pathways (Ibarra et al., 2016). Imbibed Arabidopsis seed induced into secondary dormancy by darkness showed a decline in abundance and sensitivity to GA (GA_4_; Ibarra et al., 2016). While ABA synthesis was important for the development of secondary dormancy, ABA sensitivity did not discriminate the dormancy states (Ibarra et al., 2016).

Developmentally dependent effects of exogenous GA in particular have been well-documented in buds of grapevine and other species (Lavee and May, 1997; Rinne et al., 2011; Zheng et al., 2018b, and references therein). The sensitivity of buds to exogenous ABA declined from late autumn to winter (cv. Early Sweet, 42°58’N), as the levels of endogenous ABA catabolites and *VvA8H-CYP707A4* accumulated (Zheng et al., 2015). Vines over-expressing *VvA8H-CYP707A4* showed a more rapid bud burst in explants collected from autumn through winter, relative to wild type (Zheng et al., 2018a). In a related study, exposure of explants collected in winter to GA_3_, GA_4_ or GA_7_ had an inhibitory effect on the rate of bud burst (Zheng et al., 2018b). When collected in spring, treatment of explants with GA_3_ at day 0 transiently inhibited bud burst, while treatment at day 3 accelerated it (Zheng et al., 2018b). The authors also demonstrated timing-dependent effects of exogenous GA_3_ on buds pre-treated with hydrogen cyanamide or chilling.

This knowledge provides a basis to explore the sensitivity of explants to ABA and GA collected at different developmental states and treated following periods of time in forcing conditions. For example, whether the sensitivity to GA declines in explants collected in late summer following a period of forcing conditions. Such a study may test whether the buds collected in late summer in the present study did develop a stress-induced dormancy state akin to secondary dormancy, helping to substantiate the existence of a developmental checkpoint prior to natural dormancy in the field. Notwithstanding this interpretation, the accompanying physiological and transcriptome data reflect field-state of the buds.

### ABA-dependent processes govern early seasonal changes in transcription

As described above, ABA plays important roles in the acquisition of dormancy, as well as cold, desiccation tolerance. In poplar, the function of ABA in the onset of dormancy is downstream of photoperiod responses (Tylewicz et al., 2018). In our data, the transition from summer to autumn accompanied a strong down-regulation of genes coding for the ABA biosynthesis and responses, notably the synthesis of RFOs. Additional ABA- and RFO-synthetic genes were down-regulated at the subsequent stage from autumn to winter, while ABA signalling genes were down-regulated in the final transition to spring. However, we found no clear evidence of a photoperiod response in our data. While *V. riparia* responds strongly to photoperiod, *V. vinifera* is considered a facultative long day species (Fennell and Hoover, 1991; Kühn et al., 2009; Pérez et al., 2009; Sreekantan et al., 2010; Pérez et al., 2011; Fennell et al., 2015).

ABA positively regulates the synthesis of RFOs in a number of species, which in turn play important functions in development, energy homeostasis, desiccation tolerance and response to oxidative stress (Nishizawa-Yokoi et al., 2008; Nishizawa et al., 2008; Urano et al., 2009; Sengupta et al., 2015). In glasshouse grown vines, *in situ* buds of daylength-sensitive *V. riparia* accumulated ABA content up to 21 days after exposure to short days, followed by a mild decline, while raffinose and trehalose metabolite levels did not accumulate until 28-42 days, when the buds were deemed to be dormant (Fennell et al., 2015). Four homologues of *GALACTINOL SYNTHASE* were up-regulated at the dormant stage (28-42 days) in the short day grown *V. riparia* against long day. The *V. riparia* data appear to be consistent with those seen in apple and other temperate/boreal species, where homologues of *GALACTINOL SYNTHASE* showed strong circannual rhythms, that were paralleled by the concentration of galactinol and raffinose (Cox and Stushnoff, 2001; Derory et al., 2006; Pagter et al., 2008; Ibánez et al., 2013; Falavigna et al., 2018). Wang et al. (2020) reported an increase in concentration of galactinol and stachyose in the explants following exogenous ABA. In the absence of exogenous ABA however, there was little temporal change in sugar and oligosaccharide content (Wang et al., 2020).

As described above, over-expression of *VvA8H-CYP707A4* accelerated bud burst, whether forced in autumn, winter or spring (Zheng et al., 2018a). While homologues of *A8H-CYP707A* were not differentially regulated, our data were consistent with a decline in ABA signalling responses in the transition to spring, including the down-regulation of *XERO1, DOG1, HAI2, HAI3, DRM1*, and genes coding for SNF1-related protein kinases. These data suggest that ABA and RFO levels were already high in the buds of our earliest time point and played an important function in the regulation of respiration, desiccation, acclimation to stress and physiological quiescence during the transition to autumn.

### Light and temperature responses implicate a role for blue light signalling in the resumption of growth

In our data, regulation of *CRY3, HEMA1, GUN4, PORC* and the *COR27* homologues were among the most upstream transcriptional response to light and temperature. *CRY3* encodes a cryptochrome receptor for blue and UVA light. Together with other photoreceptors, it positively regulates photomorphogenesis via the *ELONGATED HYPOCOTYL5* (*HY5*) transcription factor (Gangappa and Botto, 2016, and references therein). HY5 directly regulates other light signalling genes, including those coding for the synthesis of chlorophyll as seen prominently here; *HEMA1, GUN4, PORC* and a number of light harvesting complexes. *HY5*, together with *CRY3, HEMA1, GUN4* and *PORC* was light-regulated during bud burst (Meitha et al., 2017; Signorelli et al., 2018), but not observed in the present data. In addition, we observed down-regulation of *DRM1* and *KING1* in the transition to spring here and in light-grown buds during bud burst (Meitha et al., 2017; Signorelli et al., 2018). Both *DRM1* and *KING1* genes in Arabidopsis are repressed by blue light (Jiao et al., 2003; Kleine et al., 2007), and blue light and cytokinins function together in photomorphogenic processes during bud outgrowth (Roman et al., 2016; Signorelli et al., 2018).

The *COR27* and *COR28* homologues are key integrators of light and temperature cues in Arabidopsis (Fowler and Thomashow, 2002). The Arabidopsis COR27 and COR28 proteins are stabilised by blue light and directly interact with HY5 in transcriptional regulation of photomorphogenesis (Li et al., 2020). These proteins function downstream of CIRCADIAN CLOCK ASSOCIATED 1 (CCA1) and CONSTITUTIVE PHOTOMORPHOGENIC 1 (COP1) but upstream of floral regulators, as well as playing a role in the development of freezing tolerance (Li et al., 2016; Li et al., 2020). The prominent regulation of three *COR27* homologues from autumn through to spring is thus consistent with a regulatory function of blue light signalling via cryptochromes.

### Redox and hypoxia signalling may play important roles in structural remodelling and DNA synthesis and repair during bud development

There was prominent regulation of genes coding for redox homeostasis and carbon metabolism at each transition. Sugar and nitrogen transporters, and genes coding for gluconeogenesis were prominently down-regulated between summer and autumn, while these functions, together with activities that mobilise sugars from lipids, organic acids and oligosaccharides were up-regulated from winter to spring. In the intervening transition from autumn to winter there was little change, indicating metabolic quiescence. The synthesis of cysteine, glutathione and ascorbate was strongly regulated in the transition to spring, along with *RBOHD, ROXY1* and *ROXY2*, and other prominent redox processing functions. Moreover, ten homologues of the 49 Arabidopsis hypoxia response genes (Mustroph et al., 2009) were differentially regulated in the transition from winter to spring, together with *RAP2*.*3* and *ZPR2* (Weits et al., 2019).

Redox processing plays important metabolic roles, and ROS are intricately involved in cell expansion, cell-wall thickening and the conductivity of plasmodesmata (Gapper and Dolan, 2006; Benitez-Alfonso et al., 2011; Considine and Foyer, 2014). We have previously demonstrated strong temporal and spatial correlation between ROS and lignin abundance during bud burst in grapevine (Meitha et al., 2015), and data here indicate these processes commence well before the acute transition to bud burst. Developmental hypoxia also plays important roles in meristem functions and metabolic regulation, including during bud burst (Considine et al., 2017; Meitha et al., 2017; Gibbs et al., 2018; Weits et al., 2019; Weits et al., 2020). The pO_2_ data shown here are consistent with developmental control of hypoxia, through metabolic regulation and diffusion constraints through structural changes. It is important to indicate the strong regulation of the nuclear landscape, DNA synthesis and repair, which was most prominent in the transition towards spring. Redox processing is critical for enabling a competent rhythm during the cell cycle upon imbibition in seeds (de Simone et al., 2017), and regulated ROS synthesis is important for DNA synthesis and repair, and regulation of cell cycle transitions (Tsukagoshi et al., 2010; Velappan et al., 2017).

## CONCLUSIONS

The form and phenology of grapevine displays considerable plasticity to climate and seasonality. For the first time, we have established a field-based platform for systems approaches to study the regulation of quiescence in grapevine from bud set to bud burst. Further development of this platform will enable advanced understanding of vine phenology in a range of climate conditions. We observed an extreme resistance of explants collected in late summer to resume growth, which was not consistent with the seasonal dynamics in respiration or hydration, nor the differential gene expression. We hypothesise the existence of a physiological checkpoint following bud set, which if not met results in extreme quiescence. This reveals important considerations for interpreting bud forcing bioassays, which are widely used as a quantitative measure of dormancy. Interpretations of the field-state data however were consistent with an important regulatory role for ABA in the onset of dormancy and acclimation. Specific gene regulation of light, hypoxia and redox signalling functions during the transition to spring were accompanied by strong up-regulation of histone and chromatin regulators, together with canonical genes of the cell cycle and meristem identity. In addition, the prominent induction of aquaporin-related genes in spring suggested an important role of water transport in enabling the resumption of growth and communication prior to bud burst. Together, this study provides critical insight to the understanding of the regulation of quiescence, and prompts wider investigation of the seasonal and climate-dependencies of phenology in grapevine.

## MATERIALS AND METHODS

Unless otherwise stated, all chemicals were supplied by Sigma Aldrich, NSW, Australia.

### Plant material

Material for this study was collected from 275 similarly vigorous, non-consecutive vines of *V. vinifera* (L.) cv. Merlot (clone FVD3v14/VX/UCD on own roots), across six rows in a commercial vineyard from the Margaret River region in Western Australia, Australia (33°47’S 115°02’E). The Margaret River region has a maritime temperate climate, with mean annual temperatures 10.7-21.4 °C, predominantly winter rainfall of 957 mm per annum, and elevation 80 m (http://www.bom.gov.au/climate/averages/tables/cw_009746.shtml). Merlot is typically pruned in the first week of July, with bud burst occurring in early September. The vines used in this study were not pruned until following the final sampling in September, at which point <10 % buds had burst in the field. Vines used in this study were not treated with hydrogen cyanamide (H_2_CN_2_) in the field.

Prior to the study, the canes were tagged from numbered vines and randomly assigned to collection dates. Canes of diameter 5-12 mm, comprising nodes 2-11 acropetally (where node 1 is the node above the first internode >7 mm) were collected from specific vines from mid-summer in January through to spring in September in the years 2015 and 2016 on 12 sampling dates (28 January, 10 February, 23 February, 11 March, 23 March, 7 April, 28 April, 20 May, 7 June, 7 July, 5 August and 1 September) in 2015 and 5 sampling dates (6 January, 15 February, 10 May, 10 August and 23 September) in 2016 (Figure 1a). At the earliest sampling time (January), all basal buds up to node >15 were mature and lignified (data not shown). Immediately following sampling, the canes were stored at 20 °C in the dark for <48 h, during which time all respiratory and tissue oxygen partial pressure (pO_2_) analyses were performed. Nodes were randomly assorted to the assays, such that a single node is the basic biological unit, and where necessary, multiple buds were pooled into one biological replicate, as described for each method herein. Buds for the gene expression profiling (RNAseq) were snap-frozen at sampling and stored at −80 °C. All the material was collected between 7 to 10 am, to avoid changes in gene expression due to circadian regulation.

### Depth of dormancy (BB_50_)

Bud dormancy was measured using single node explants, which is a commonly used system to study the physiology of early shoot and inflorescence development in grapevine (Pouget, 1963; Mullins, 1966; Nigond, 1967; Antolín et al., 2010), and excludes the influence of adjacent or distant organs. Each explant comprised *ca*. 50 mm of cane beneath and 10 mm cane above the node, diameter *ca*. 7-12 mm. Explants were grown in potting mix (pH∼6.0 fine composted pine bark: coco peat: brown river sand, ratio 2.5:1:1.5 (w/w)) in a controlled temperature room at 20 °C, 12 h photoperiod, illuminated with fluorescent light at 100 µmol.m^-2^.s^-1^. Soil moisture was maintained at >80% of water holding capacity.

Because bud burst varies stochastically, 50 buds were used to represent the dormancy state of the population at the time of sampling. H_2_CN_2_ was used as a positive control, as it is a widely used dormancy breaking agent and has also been widely studied in the context of bud dormancy. For the H_2_CN_2_ treatment, 50 buds were immersed in 2.5 % v/v Dormex® (Crop Care, Australasia; equal to 0.31 M H_2_CN_2_), for 20 seconds and air dried prior to transplanting. The untreated buds were immersed in distilled water for 20 s and air dried prior to planting. The stage of bud burst was scored at the emergence of visible green leaf tips (EL4), according to the modified Eichorn-Lorenz scale (Coombe, 1995). Bud burst was recorded three times per week for up to 350 d or until 100 % bud burst, and the depth of dormancy was calculated as the time to reach 50 % bud burst (25/50 buds burst; BB_50_).

### Climate and chilling data

The weather data was obtained from the Moss Wood vineyard in Margaret River, Western Australia for 2015 and 2016. Average daily mean air temperatures were calculated from hourly air temperatures using the R statistical package (R Core Team, 2020) and plotted using ggplot2 package of R (Wickham, 2009) as scatter dot plots fitted with a quadratic spline with degree of freedom (df)=4 and degree=2. The photoperiod data was obtained from the website http://aa.usno.navy.mil/cgi-bin/aa_rstablew.pl and plotted using the ggplot2 package of R (Wickham, 2009) as a line plot.

Chilling data were modelled by three methods; the Daily Positive Utah Chill Unit (DPCU) model (Linsley-Noakes and Allan, 1994), the Utah model (Richardson et al., 1974), and the base 7.2 °C model. The cumulative chilling was calculated for each day from the raw data and represented as a line plot using R package (R Core Team, 2020) and ggplot2 package of R (Wickham, 2009) respectively.

### Bud moisture content

Ten buds per biological replicate (3 biological replicates) were transversely sectioned from the canes and their fresh weight was recorded. All buds were inspected for signs of necrosis prior to including them in the analysis. Dry weight was calculated post-drying at 60 °C for 7 d, and moisture content was calculated as g H_2_O.100g^-1^ fresh weight.

### Bud respiration

Five buds per biological replicate (3 biological replicates) were transversely sectioned from the canes and weighed immediately and placed on a thin agar plate, sectioned side down, to prevent dehydration and gas exchange from the cut base. Rate of O_2_ uptake and CO_2_ release were calculated.

*O*_*2*_ *uptake*: The rate of O_2_ uptake for every biological replicate was measured in the dark using Unisense MicroRespiration system (OX-MR, Unisense, Denmark) with Clark-type O_2_ microsensor in a 4 mL respiration chamber at a constant temperature of 20 °C (Shaw et al., 2017). The readings were obtained using the SensorTrace RATE software (Unisense, Denmark).

*CO*_*2*_ *release*: The rate of CO_2_ release was measured in the dark, in an insect respiration chamber (6400-89; Li-COR, Lincoln, NB, USA) attached to a Li-6400XT portable gas exchange system at 20 °C, in CO_2_-controlled air (380 µmol CO_2_.mol^-1^ air) with 100 µmol.m^-1^.s^-1^ air flow, at 55–75 % relative humidity. The measurements were recorded once the ‘stableF’ value read 1 (i.e. after stabilization of humidity, CO_2_ and air flow) following transfer of sample to the chamber. The readings obtained were later analysed.

### Internal bud O_2_ partial pressure (pO_2_)

The internal pO_2_ of 3 to 4 in 2015 (6 in 2016) biological replicates with one single bud cutting per replicate was measured in the dark at 20 °C using a Clark-type oxygen micro-sensor with a tip diameter of 25 µm (OX-25; Unisense A/S, Aarhus, Denmark). The microelectrode was calibrated at atmospheric pO_2_ of 20.87 kPa and at zero pO_2_ (100 % nitrogen gas). Then mechanically guided into the bud, starting from the outer scale surface at 0 µm to the inner meristematic core at 2000 µm, in 25 µm steps with a stabilizing pause of 3 s in between steps with the aid of a motorized micro-manipulator, as previously described (Shaw et al., 2017). The values were recorded automatically at the end of each step (i.e. every 25 µm). The readings were processed using the SensorTrace RATE software (Unisense, Denmark), analysed using R statistical package (R Core Team, 2020) and presented in graphical form using the ggplot2 package of R (Wickham, 2009), data fitted with a LOESS regression curve at 95 % confidence intervals (n=3 to 6 per month per year).

### Physiological data analysis and statistics

All calculations were performed using Microsoft Excel 2016 and R statistical package (R Core Team, 2020) and graphics were compiled using the ggplot2 package of R (Wickham, 2009). At least three biological replicates were used per analysis. Significant differences among various sampling dates were corroborated statistically by applying one-way ANOVA test, using Tukey’s honestly significant difference (HSD) posthoc test with *P*≤0.01.

### RNA isolation, library preparation and RNAseq

RNA extractions and libraries (three biological replicates of 2-3 buds each per condition, *ca*. 50 ng) were prepared from material collected at the corresponding dates (Figure 1a) and prepared for sequenced as described in Meitha et al. (2017), with minor modifications. Libraries were prepared with the TruSeq Stranded Total RNA with Ribo-Zero Plant Kit (Illumina, Scorseby, Australia) according to manufacturer’s instructions. RNAseq was performed on an Illumina HiSeq2500 by Novogene (Hong Kong) at *ca*. 9 Gb data of 150 pair-end (PE) reads per library. Raw data files have been submitted to NCBI (BioProject ID PRJNA575976, http://www.ncbi.nlm.nih.gov/bioproject/575796).

### RNAseq data processing and analysis

Transcriptomic data analysis was performed according to Meitha et al. (2017) with minor modifications, and summary statistics of read length and read mapping are provided in Supplemental Table S2. Briefly, FastQC software was used to assess the quality of the fastq files (https://www.bioinformatics.babraham.ac.uk/projects/fastqc/). Adapter and quality trimming were performed using Trimmomatic (Bolger et al., 2014) with default settings. Salmon (Patro et al., 2017), a pseudoalignment algorithm was used to map the trimmed reads to the 12X v2.1 *V. vinifera* PN40024 reference genome (Canaguier et al., 2017) with sequence-specific and GC bias correction, and quantify gene expression. The grapevine reference genome and annotation were obtained from the Phytozome v.12.1 database (Goodstein et al., 2012). The counts matrix obtained from Salmon was read into edgeR (Robinson et al., 2010), then normalized using the trimmed mean of M values (TMM) method to log counts per million reads (logCPM) and filtered to remove lowly expressed genes i.e. genes with <1 reads in any sample. Graphical representations of the data before and after normalization are presented in Supplemental Figure S2 (a and b). The quality of the replicates was checked using unsupervised clustering of samples before and after normalisation and presented in the multi-dimensional scaling (MDS) plot (Supplemental Figure S2c). The normalized counts for each gene were further transformed and fitted into a linear model using the Voom function from limma (Law et al., 2014). Mean-variance relationships were evaluated before and after Voom precision weights were applied to the data (Supplemental Figure S3).

Differentially expressed gene (DEG) analysis was carried out using edgeR and limma (Robinson et al., 2010; Ritchie et al., 2015) Bioconductor packages with default settings. *P* values were corrected for multiple testing using the Benjamini-Hochberg method (FDR ≤ 0.01) (Benjamini and Hochberg, 1995). The data were then filtered to consider only genes with an absolute Log_2_Fold Change ≥ 1 (Log_2_FC |1|).

### Functional enrichment analysis

Three lists of DEGs from each comparison, up- and down-regulated genes and all (up-plus down-regulated genes) were used to perform a functional enrichment analysis using GOstats (Falcon and Gentleman, 2007) in R. This was done to identify significant functional categories of the genes based on *V. vinifera* functional classification of 12X v. 2.1. GOstats uses the hypergeometric distribution (a widely used method to test for overrepresentation), to compare each DEG list with the list of total genes. Gene ontology (GO) terms with *P*<0.05 were considered to be significantly enriched. All plots were rendered using the gplots and ggplots programs in R.

### Known *cis*-regulatory motif analysis

Grapevine promoter sequences (1.2 kb upstream of the coding sequence) of all annotated *V. vinifera* 12X v.2.1 genes (which is referred to as the background) were obtained from the Grape Genome Database (Vitulo et al., 2014). Subsets of promoter sequences of DEGs from the three comparisons February-April, April-June, June-September were retrieved from all Grapevine genes promoter sequences i.e. the background. Each gene set was analysed for enriched transcription factor binding motifs/cis-elements using by using Homer v.4.11 (Heinz et al., 2010) findMotifs.pl script with default parameters.

## Acknowledgements

We are extremely grateful to Keith Mugford and his team of Moss Wood Vineyards in Margaret River for their generous information, support and for enabling the sampling to continue at irregular times. We also acknowledge support and teamwork of other laboratory members, particularly Dr Karlia Meitha, Dr Dina Hermawaty, Juwita Dewi and Wisam Salo.

## Supplemental data files

**Supplemental Figure S1. Seasonal changes in photoperiod, temperature and cumulative chilling in the Margaret River region of Western Australia (2015). (a)** Daylength (Source: https://aa.usno.navy.mil), **(b)** minimum (grey) and maximum (black) daily air temperature: local, daily, uncorrected records), and **(c)** cumulative chill units by three models: Daily Positive Utah Chilling Units (Linsley-Noakes and Allan, 1994), Utah model (Richardson *et al*., 1974) and base 7.2 °C. All values were calculated from the raw data collected at 15 min intervals.

**Supplemental Figure S2**. Box plots of read counts/effective library sizes before and after normalization and a multi-dimensional scaling (MDS) plot showing the similarities and dissimilarities between samples. Boxplots of log2 values showing expression distributions **(a)** before and **(b)** after normalization. **(c)** A multi-dimensional scaling (MDS) plot showing the similarities and dissimilarities between samples after normalization

**Supplemental Figure S3**. Scatterplots of the distribution of means (x-axis) and variances (y-axis) of each gene showing the dependence between the two **(a)** before and **(b)** after Voom is applied to the data. Voom extracts residual variances from fitting linear models to log-CPM transformed data.

**Supplemental Table S1**. Bud burst, physiological and climate data for 2015 and 2016 Merlot buds grown in Margaret River region of Western Australia (33°S latitude). **(a)** Bud burst, days to 50 % bud burst (n=50). **(b)** Respiration and hydration (n=3). **(c)** Chilling data according to the Positive Daily Chilling Units model. **(d)** Rainfall.

**Supplemental Table S2**. RNA-Seq data statistics describing reads pre- and post-quality control and number of reads that mapped to the genome (effective library size).

**Supplemental Table S3**. Number of genes differentially expressed (FDR ≤ 0.01, log2FC |1|) in developmental comparisons from grapevine buds sampled from summer to spring in the Margaret River region of Western Australia in 2015. Comparisons are developmental transition for April/February, June/April and Sept/June sampling dates (NCBI BioProject ID PRJNA575976, http://www.ncbi.nlm.nih.gov/bioproject/575796).

**Supplemental Table S4**. Statistically significant gene ontology (GO) terms of differentially expressed genes in grapevine buds sampled from summer to spring in the Margaret River region of Western Australia in 2015. Comparisons are developmental contrasts for April/February, June/April and Sept/June sampling dates.

**Supplemental Table S5**. A subset of Supplemental Table S3 referring to specific genes outlined in the manuscript. Comparisons are developmental transition for April/February, June/April and Sept/June sampling dates. An additional worksheet shows a comparison of differentially expressed genes in the present study with that from Meitha et al., 2017 and

Signorelli et al., 2018; data are conditionally formatted for up-regulated genes (green) and down-regulated genes (blue).

**Supplemental Table S6**. Statistically enriched known *cis*-elements/motifs 1.2 kB upstream the transcriptional start site (TSS) of differentially expressed genes in grapevine buds sampled from summer to spring in the Margaret River region of Western Australia in 2015. Comparisons are developmental contrasts for April/February, June/April and Sept/June sampling dates.

**Supplemental Table S7**. Homologues of cell cycle, aquaporin and AP2/ERF genes and their expression patterns in the DEG data set, as used for Figure 5.

